# Transferrin receptor binding valency determines the circulating half-life and weight-loss efficacy of Fc-FGF21 in TfR-humanized mice

**DOI:** 10.64898/2026.11.02.763770

**Authors:** Hong Zhou, Yanru Deng, Huiqiao Li, Jieyi Meng, Leon G. Straub, Shangang Zhao, Yi Zhu

**Affiliations:** USDA/ARS Children’s Nutrition Research Center, Department of Pediatrics, Baylor College of Medicine, Houston, TX, USA; Department of Pharmacology, University of Virginia School of Medicine, Charlottesville, VA, USA; Robert M. Berne Cardiovascular Research Center, Charlottesville, VA, USA; Barshop Institute for Longevity and Aging Studies, Division of Endocrinology, Department of Medicine, University of Texas Health Science Center at San Antonio, San Antonio, TX, USA

**Keywords:** fibroblast growth factor 21, transferrin receptor, receptor binding valency, blood–brain barrier, circulating half-life, weight loss, Fc fusion protein

## Abstract

Fibroblast growth factor 21 (FGF21) is a promising metabolic hormone with potent anti-obesity effects. It acts in both the central nervous system and peripheral tissues to exert its metabolic benefits. Because its receptors are expressed at low levels in the brain, we hypothesized that enhanced transport across the blood-brain barrier (BBB) would improve its efficacy. To achieve this, FGF21 was fused to Fc(TV35), an engineered Fc fragment that binds the human transferrin receptor (hTfR), to generate Fc(TV35)-FGF21. Fc(TV35)-FGF21 does not bind to mouse TfR, so it produced weight-loss effects in wild-type diet-induced obese mice comparable to those of Fc-FGF21. However, it failed to induce weight loss in homozygous hTfR-humanized mice because of a markedly shortened circulating half-life. Converting Fc(TV35)-FGF21 from a homodimer to a bispecific Fc format with a single hTfR-binding arm partially restored the circulating half-life and weight-loss efficacy. Reducing mouse hTfR zygosity from homozygous to heterozygous did not substantially improve the engineered protein’s half-life or weight-loss effect. Taken together, our findings suggest that sustained systemic exposure is a major determinant of FGF21’s weight-loss efficacy and that hTfR engagement can shorten its systemic half-life and compromise its therapeutic activity. These results highlight an important design limitation of TfR-based brain vehicles, especially for delivering metabolic hormones with low-abundance CNS receptor targets. As a result, further optimization of shuttle formats to minimize systemic clearance while crossing the BBB is needed for metabolic hormones.

## Introduction

Metabolic hormones such as insulin, leptin, glucagon-like peptide-1 (GLP-1), and fibroblast growth factor 21 (FGF21) are critical regulators of systemic energy and glucose homeostasis, acting on both peripheral tissues and the central nervous system (CNS) to coordinate metabolic responses to nutritional status [1-7]. Accumulating evidence indicates that the CNS contributes substantially to the metabolic actions of these hormones, influencing appetite, energy expenditure, and thermogenesis [1-4]. However, the CNS exposure of circulating peptide hormones is limited and tightly regulated by the blood–brain barrier (BBB), a tightly regulated endothelial interface that restricts the entry of macromolecules into the brain parenchyma [8]. Thus, strategies to increase CNS exposure to metabolic hormones may enhance their efficacy in treating obesity and metabolic disorders.

The transferrin receptor 1 (TfR1) is abundantly expressed on the luminal surface of brain endothelial cells, facilitates receptor-mediated transcytosis across the BBB [9], and has been widely exploited as a molecular “shuttle” to deliver therapeutic cargo into the CNS [10]. Engineered antibodies and Fc fragments with optimized TfR-binding properties have been successfully used to achieve enhanced brain exposure [11]. This approach has been applied to deliver enzymes [12], antibodies [13], and neurotrophic factors [14] for the treatment of neurological diseases.

In the present study, we generated an engineered Fc–FGF21 fusion protein in which the Fc domain was further modified to bind human, but not mouse, TfR. By enabling receptor-mediated transcytosis, we aimed to increase the CNS exposure of FGF21 and thereby enhance its body weight-lowering efficacy. We evaluated this protein in TfR-humanized diet-induced obese (DIO) mice to determine whether this hTfR1-targeted brain design would potentiate its weight-reducing and metabolic effects compared with those of conventional Fc-FGF21 analogs.

## Methods

### Mouse model and treatment

All animal protocols were approved by the Institutional Animal Care and Use Committee of Baylor College of Medicine. hTfR1^homo^ mice (cat. no. NM-HU-215059) were purchased from Shanghai Model Organisms. WT C57BL/6J mice were obtained from The Jackson Laboratory. hTfR1^het^ mice were generated by crossing hTfR1^homo^ mice with WT C57BL/6J mice. Mice were housed under a 12-h light/12-h dark cycle in a temperature-controlled environment (22°C), with ad libitum access to food and water, and under routine veterinary supervision. Mice were maintained on a standard rodent chow diet (Bio-Serv, cat. no. AIN-76A) or a high-fat diet (HFD; Bio-Serv, cat. no. S1850). Mice were treated with fusion protein at 9.1 nmol/kg body weight (1 mg/kg for Fc-FGF21 protein) via i.p. injection, three times per week. Animals were monitored throughout treatment for body-weight changes and general condition, including activity, posture, grooming, hydration, and signs of injection-related distress. Despite the substantial treatment-induced weight loss, no overt adverse clinical signs, treatment-related deaths, or humane-endpoint removals were observed. Only male mice were used in the experiments. Mice were housed individually for food-intake measurements.

### Protein design and purification

To generate the fusion proteins, the N-terminus of FGF21 was engineered onto the Fc domain via a flexible linker. Plasmid constructs encoding the various FGF21 and Fc protein variants, as well as procedures for plasmid transformation, protein expression, and purification using a bacterial system, were described previously [11, 15, 16]. However, because the bacterial expression system failed to yield the target proteins, the Expi293F mammalian cell expression system was subsequently used for protein production. Briefly, Expi293F cells were transfected with plasmids using PEI MAX and cultured for 3 days. The culture supernatants were collected, and the recombinant proteins were purified by Ni-affinity chromatography, eluted with imidazole, and dialyzed overnight against PBS, following standard procedures. Protein purity was assessed by SDS–PAGE under reducing and non-reducing conditions. Only preparations with >95% purity were used in the experiments. All Fc–FGF21 fusion proteins in this experiment were produced by Synbio Technologies (Monmouth Junction, NJ, USA).

### Measurement of the circulating half-life of Fc-FGF21

To determine the circulating half-lives of the recombinant proteins *in vivo*, mice were injected via the tail vein with the indicated fusion protein at 9.1 nmol/kg body weight. At the indicated time points, blood was collected from the tail tip and centrifuged to isolate serum. Serum concentrations of the fusion proteins were quantified using a human IgG1 ELISA kit (ThermoFisher, cat. no. BMS2092) according to the manufacturer’s protocol, except that the engineered Fc-FGF21 molecular weight was used to calculate final concentration. The apparent terminal half-life was estimated by log-linear regression of the natural log-transformed serum concentrations at the 8-, 24-, and 48-h time points. For Fc(TV35)-FGF21, the 4-, 8-, and 24-h time points were selected because its concentrations at 24 h and 48 h were extremely low and comparable.

### Measurement of whole-brain FGF21 levels

At 46 h after intraperitoneal injection of Fc–FGF21 or Fc(TV35)–FGF21, blood samples were collected from the tail vein. The mice were then anesthetized and transcardially perfused with 25–30 mL of ice-cold PBS to remove residual blood from the cerebral vasculature. Whole brains were rapidly collected, snap-frozen in liquid nitrogen, and stored at −80°C. A piece of brain tissue was then homogenized in T-PER^™^ Tissue Protein Extraction Buffer (ThermoFisher, cat. no. 78510) supplemented with a protease inhibitor cocktail (Fisher Scientific, cat. no. 78442) using TissueLyser. The homogenates were subsequently centrifuged at 10,000 × g for 5 min at 4°C. The resulting supernatants were collected, and Fc–FGF21 concentrations were quantified using the human IgG1 ELISA kit to detect human IgG1 Fc.

### Data analysis

Data are presented as the mean□±□standard error of the mean (SEM). Statistical analyses were performed using GraphPad Prism software (version 10). n represents the number of animals per experimental group. An unpaired, two-tailed Student’s *t-*test was used to compare two independent groups. For mouse experiments, each biological replicate corresponded to a separate mouse. Differences among multiple conditions and between groups over time were evaluated using two-way ANOVA. Statistical significance was defined as *p* < 0.05. The statistical test used for each dataset is specified in the corresponding figure legend.

## Results

### Design and purification of Fc(TV35)-FGF21

Given that native FGF21 has moderate efficacy for weight loss in DIO mice because of its short half-life [17], we fused FGF21 to the C-terminus of Fc to extend its half-life using a published strategy [15]. Several mutations (L98R, P171G, and A180E) were incorporated into FGF21 to further extend its half-life and reduce aggregation and *in vivo* degradation [15]. To enhance brain delivery, mutant FGF21 was fused to an engineered Fc-based transport vehicle (TV) designed to bind human transferrin receptor 1 (hTfR1, encoded by TFRC) [11], and the resulting construct was named Fc(TV35)-FGF21 (Fig. 1A). The TV acts as a “Trojan horse” by binding to the apical domain of hTfR1 at the BBB, thereby facilitating receptor-mediated transcytosis across brain endothelial cells [11]. Among the engineered TV variants, TV35 was selected for fusion with mutant FGF21 because its moderate hTfR1-binding properties are tuned for efficient BBB transport [11]. In the resulting construct, TV35 is presented in a bivalent format through the dimeric Fc scaffold (Fig. 1A).

**Figure 1.**
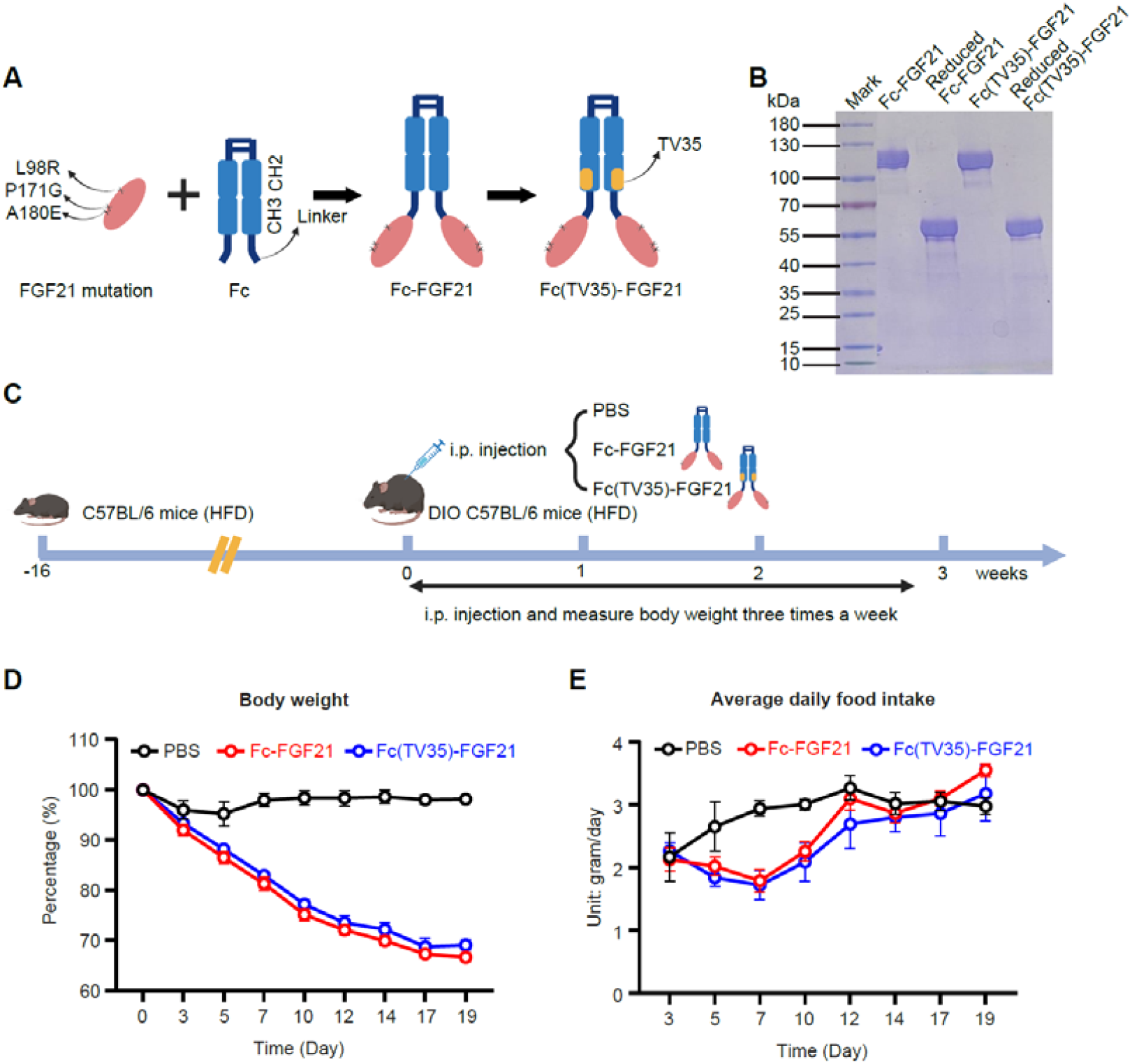
Design, purification, and function of Fc(TV35)-FGF21. (A) Schematic diagrams of the Fc-FGF21 and Fc(TV35)-FGF21 constructs. (B) SDS-PAGE analysis of purified Fc-FGF21 and Fc(TV35)-FGF21. (C) Schematic illustration of the experimental design for the studies shown in panels D and E. C57BL/6J mice were fed a high-fat diet (HFD) for 16 weeks to establish diet-induced obesity (DIO). DIO mice were then treated with PBS, Fc-FGF21, or Fc(TV35)-FGF21 three times per week, and body weight and food intake were monitored throughout the study. (D) Relative body weight over time, presented as a percentage of body weight on day 0. DIO mice maintained on an HFD were treated with PBS, Fc–FGF21, or Fc(TV35)–FGF21 at the indicated times (n = 8 for each group). (E) Average daily food intake calculated three times per week (n = 8 for each group). All data are presented as the mean ± s.e.m. *P* values were calculated using a mixed-effects two-way ANOVA.

We first attempted to produce the two target proteins, Fc-FGF21 and Fc(TV35)-FGF21, using a previously reported bacterial expression system and denaturation/refolding purification methods [16]. However, this approach failed to yield the desired proteins, probably because of improper folding. We therefore switched to the Expi293F mammalian expression system. SDS-PAGE analysis showed that both target proteins were successfully produced (Fig. 1B and Fig. S1). Because Fc(TV35) does not bind mouse TfR (mTfR), we tested the weight-loss efficacy of Fc(TV35)-FGF21 in DIO C57BL/6J mice without hTfR expression (Fig. 1C). There was no significant difference in weight loss between mice treated with Fc-FGF21 and those treated with Fc(TV35)-FGF21 (Fig. 1D), suggesting that introduction of the TV35 mutation into the Fc region did not affect FGF21 efficacy, as expected. Food intake was transiently suppressed after treatment with either Fc-FGF21 or Fc(TV35)-FGF21 and then returned to levels observed in PBS-treated mice. After 12 days, there was no difference in food intake between the Fc-FGF21 and Fc(TV35)-FGF21 groups (Fig. 1E).

### TV35-mediated bivalent hTfR binding completely abolished the weight-loss effect of FGF21 in hTfR^homo^ mice

Although mTfR1 and hTfR1 are highly homologous, their binding regions differ substantially, particularly in the apical domain. Consequently, antibodies against one ortholog generally do not cross-react with the other. TV35 is engineered by directed evolution of an Fc domain to selectively bind the apical domain of hTfR1. It exhibits species-dependent TfR1 binding, with affinities of approximately 100 nM for hTfR1 and ∼2 μM for cynomolgus monkey TfR1, while showing no detectable reactivity toward mTfR1 [11]. To better evaluate the effect of TV35-mediated bivalent hTfR binding on FGF21-induced weight loss, we used fully humanized hTfR1 mice. Homozygous hTfR (hTfR^homo^) mice were viable and fertile. hTfR^homo^ mice appeared normal but were moderately resistant to high-fat diet (HFD)-induced weight gain, based on a comparison of their body-weight trajectory with data from a historical wild-type (WT) C57BL/6J mouse cohort (Fig. S2). TfR is essential for erythropoiesis and nervous-system development, and germline homozygous knockout of TfR results in early embryonic death [18]. The slight resistance of hTfR^homo^ mice to HFD-induced obesity may reflect a mild functional iron-restricted state caused by a species mismatch between mouse transferrin and the humanized receptor, creating a mild, chronic “cellular iron deficiency” that directly affects adipose tissue expansion and increases energy expenditure [19, 20].

To compare the weight-loss effects of Fc-FGF21 and Fc(TV35)-FGF21, hTfR^homo^ mice were fed an HFD for 14 weeks to induce obesity. Then, mice were administered Fc-FGF21 or Fc(TV35)-FGF21 by intraperitoneal injection three times a week (Fig. 2A). To our surprise, introduction of TV35-mediated bivalent hTfR binding completely abolished the weight-loss effect of FGF21 in hTfR^homo^ mice (Fig. 2B). This effect coincided with a significantly shorter circulating half-life of Fc(TV35)-FGF21 (4.3 ± 0.8 h for Fc(TV35)-FGF21 versus 50.9 ± 9.0 h for Fc-FGF21; Fig. 2C), significantly lower serum protein levels at 20 and 46 h after i.p. injection (Fig. 2D), and lower brain FGF21 protein content at the end of the experiment (46 h; Fig. 2E).

**Figure 2.**
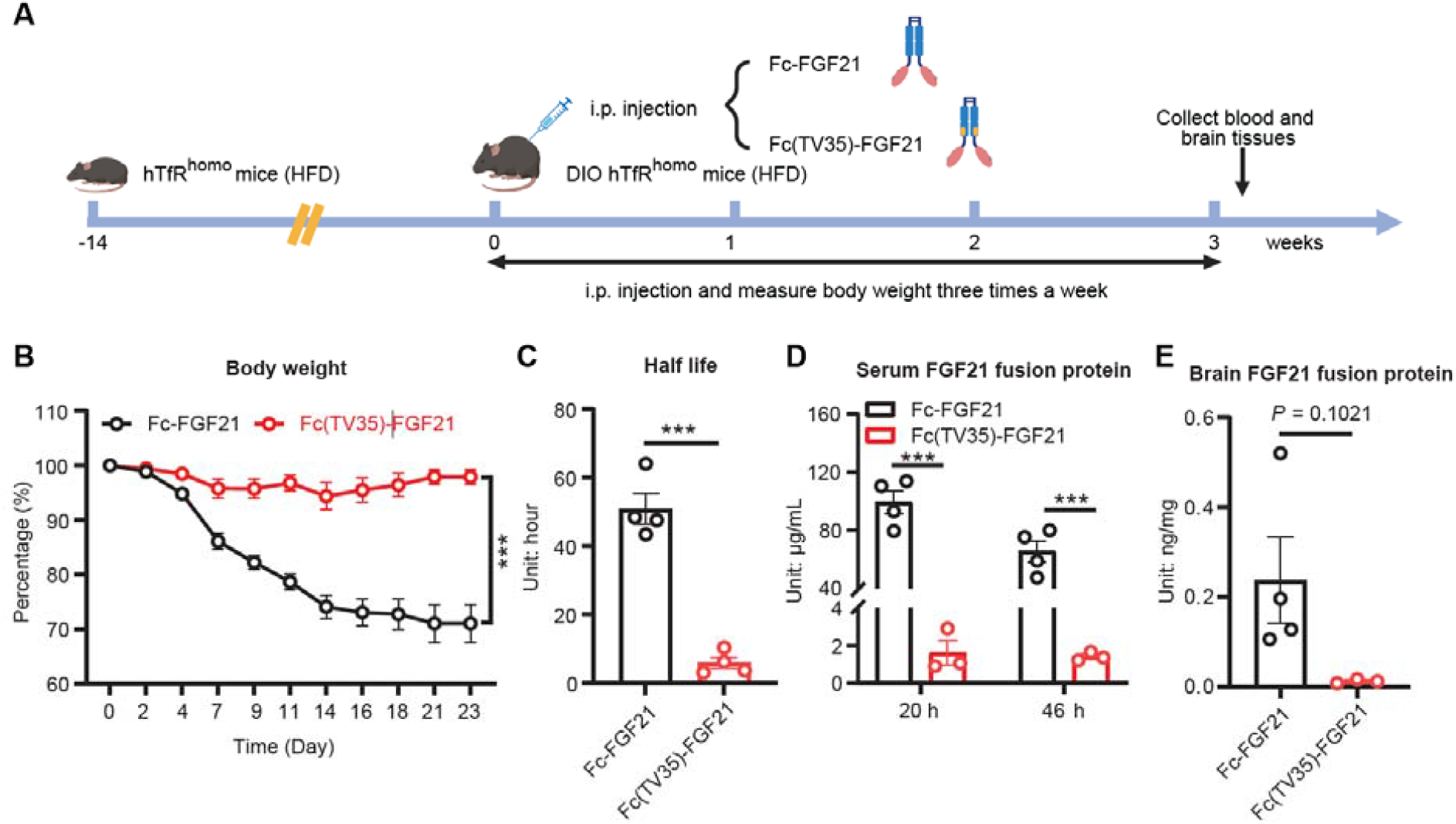
TV35-mediated bivalent hTfR binding shortens Fc-FGF21’s serum half-life and abolishes its weight loss in hTfR^homo^ mice. (A) Schematic diagram of the experimental design for hTfR^homo^ mice treated with Fc-FGF21 or Fc(TV35)-FGF21. (B) Relative body weight over time, presented as a percentage of body weight on day 0. DIO hTfR^homo^ mice were treated with Fc–FGF21 or Fc(TV35)–FGF21 three times per week (n = 4 for the Fc–FGF21 group; n = 3 for the Fc(TV35)–FGF21 group). (C) Serum half-lives of Fc-FGF21 and Fc(TV35)-FGF21 in hTfR^homo^ mice (n = 4 per group). (D) Serum Fc–FGF21 or Fc(TV35)–FGF21 levels in hTfR^homo^ mice at 20 and 46 h after the final injection on day 21 (n = 4 for the Fc–FGF21 group; n = 3 for the Fc(TV35)–FGF21 group). (E) Brain Fc–FGF21 or Fc(TV35)–FGF21 levels in hTfR^homo^ mice at 46 h after the final injection on day 21 (n = 4 for the Fc–FGF21 group; n = 3 for the Fc(TV35)– FGF21 group). All data are presented as the mean ± s.e.m. \*\*\**p* < 0.001, as determined using a mixed-effects two-way ANOVA or Student’s *t*-test, as appropriate.

### Monovalent hTfR binding partially restored the weight-reducing efficacy of Fc(TV35)-FGF21

We reasoned that the bivalent TV35 construct might undergo predominant intracellular degradation of the Fc-FGF21 fusion protein rather than transport it across the endothelial layer. To avoid bivalent hTfR1 engagement, we used the knob-into-hole heterodimerization strategy to generate monovalent hTfR1-binding Fc-FGF21 fusion proteins. This strategy utilizes preferential pairing of two distinct Fc chains through complementary mutations in the Fc CH3 domain. One heavy chain carries bulky “knob” mutations (S354C and T366W), while the other contains complementary “hole” mutations formed by smaller residues (Y349C, T366S, L368A, and Y407V) [21, 22]. These mutations create steric complementarity that favors heterodimer formation over homodimerization. Using this approach, we generated three heterodimeric Fc-FGF21 proteins (Fig. 3A): a TV35-free Fc-FGF21 variant (referred to as Fc(knob-hole)-FGF21) as the control, and two monovalent Fc(TV35)-FGF21 variants. Depending on whether TV35 was placed on either the knob Fc chain or the hole Fc chain, they are named Fc(knob-TV35)-FGF21 and Fc(hole-TV35)-FGF21.

**Figure 3.**
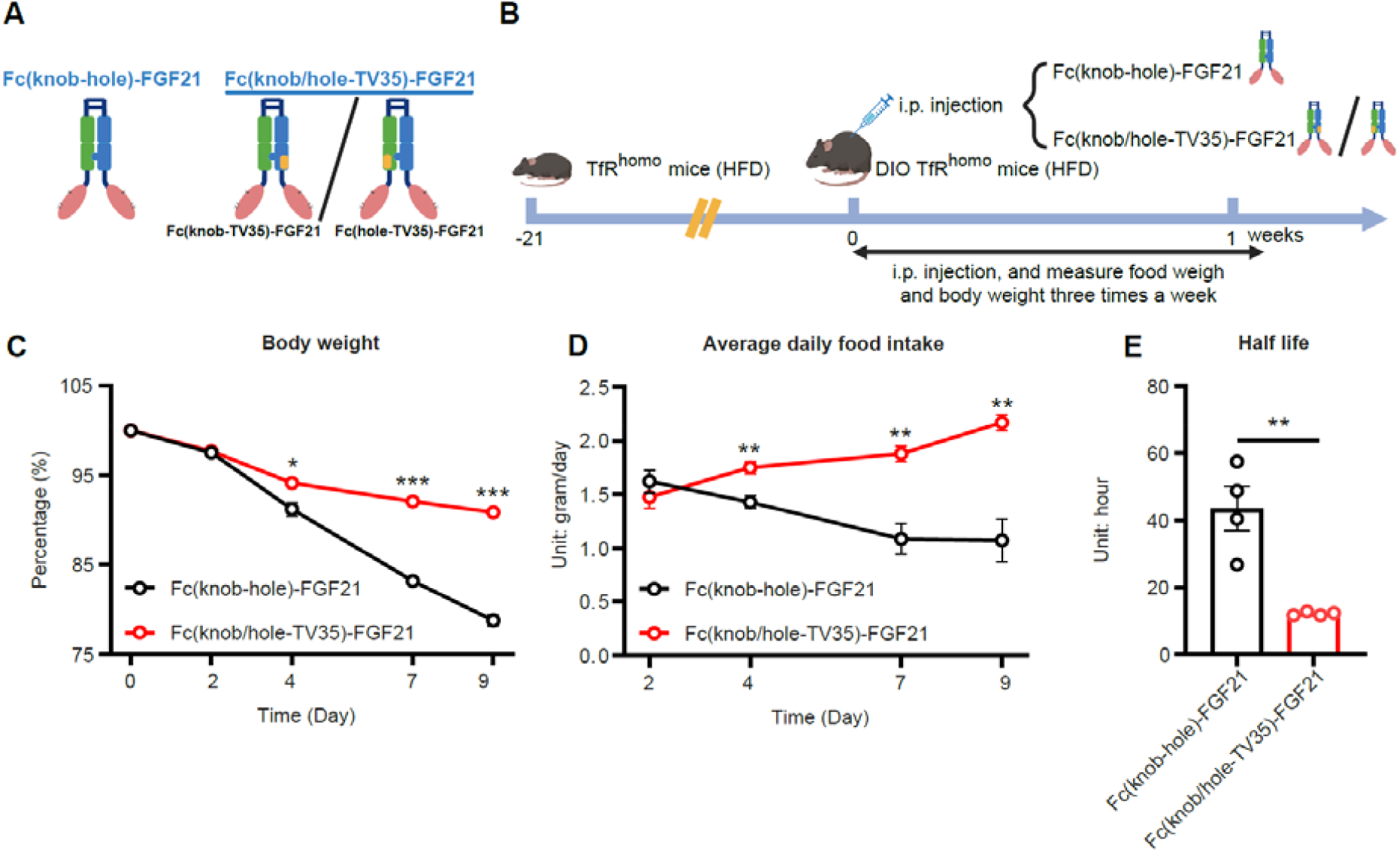
TV35-mediated monovalent hTfR binding still shortened the serum half-life but partially restored the weight-loss efficacy of Fc(TV35)-FGF21 in hTfR^homo^ mice. (A) Schematic diagrams of the bispecific Fc-FGF21 fusion proteins, Fc(knob-hole)-FGF21 and Fc(knob/hole-TV35)-FGF21 (including Fc(knob-TV35)-FGF21 and Fc(hole-TV35)-FGF21). (B) Schematic diagram of the mouse treatment design. (C) Relative body weight over time, presented as a percentage of body weight on day 0. DIO hTfR^homo^ mice were treated with Fc(knob-hole)-FGF21 or Fc(knob/hole-TV35)-FGF21 three times per week (n = 7 for the Fc(knob-hole)-FGF21 group; n = 10 for the Fc(knob/hole-TV35)-FGF21 group, including n = 5 for Fc(knob-TV35)-FGF21 and n = 5 for Fc(hole-TV35)-FGF21). (D) Corresponding average daily food intake per mouse, calculated over the specified intervals (days 0–2, 2–4, 4–7, and 7–9). (E) Serum half-lives of Fc(knob-hole)-FGF21 and Fc(knob/hole-TV35)-FGF21 in hTfR^homo^ mice (n = 4 for the Fc(knob-hole)-FGF21 group; n = 4 for the Fc(knob/hole-TV35)-FGF21 group, including n = 2 for Fc(knob-TV35)-FGF21 and n = 2 for Fc(hole-TV35)-FGF21). All data are presented as the mean ± s.e.m. \**p* < 0.05, \*\**p* < 0.01, and \*\*\**p* < 0.001, as determined using a mixed-effects two-way ANOVA or Student’s *t-*test, as appropriate.

We next examined whether this updated engineering strategy would enhance the weight-loss efficacy of FGF21 *in vivo*. To test this, DIO hTfR^homo^ mice were administered different engineered FGF21 proteins (Fig. 3B). As expected, the position of TV35 on the Fc chain did not affect its efficacy, supported by the fact that there were no statistically significant differences in weight loss, food intake, and half-life between Fc(knob-TV35)-FGF21 and Fc(hole-TV35)-FGF21 in hTfR^homo^ mice (Fig. S3A–C). Because both constructs were designed to provide monovalent hTfR1 binding, data from mice treated with either configuration were pooled for the subsequent valency-based analyses and are referred to collectively as Fc(knob/hole-TV35)-FGF21. Body weight decreased by 9.1% in the Fc(knob/hole-TV35)-FGF21 group on day 9 (Fig. 3C), compared with the completely abolished weight-loss effect of Fc(TV35)-FGF21, suggesting that the updated engineering strategy was partially successful in restoring Fc-FGF21’s weight-loss effect. The weight-loss effect was accompanied by reduced food intake (Fig. 3D). In addition, the circulating half-life of Fc(knob/hole-TV35)-FGF21 was 12.1 ± 0.5 h, 2.8-fold longer than that of Fc(TV35)-FGF21 but still markedly shorter than the 43.4 ± 13.1 h half-life of Fc(knob/hole)-FGF21 (Fig. 3E). Altogether, these data suggest that monovalent hTfR binding still leads to faster systemic elimination of the Fc-FGF21 protein and reduces Fc-FGF21 weight-loss efficacy.

### Reducing hTfR gene dosage did not improve the weight-loss efficacy of monovalent Fc(knob/hole-TV35)-FGF21

Encouraged by the partial recovery of the weight-loss effect achieved by reducing the valency of Fc(TV35)-FGF21, we asked whether heterozygous hTfR (hTfR^het^) mice would allow the monovalent Fc(knob/hole-TV35)-FGF21 protein to achieve a greater weight-loss effect. To test this, DIO hTfR^het^ mice were administered Fc(knob/hole)-FGF21 or Fc(knob/hole-TV35)-FGF21, and body weight and food intake were monitored (Fig. 4A). Fc(knob/hole-TV35)-FGF21 was less effective than Fc(knob/hole)-FGF21 in inducing weight loss and suppressing food intake in hTfR^het^ mice (Fig. 4B and C). Of note, there was no significant difference in weight loss effects of Fc(knob/hole-TV35)-FGF21 between hTfR^het^ mice and hTfR^homo^ mice in the cross-cohort comparison (Fig. 4D). The circulating half-life of Fc(knob/hole-TV35)-FGF21 in hTfR^het^ mice was 15.5 ± 1.1 h (Fig. 4E), significantly shorter than that of Fc(knob/hole)-FGF21, as seen in hTfR^homo^ mice. As expected, Fc(knob-TV35)-FGF21 and Fc(hole-TV35)-FGF21 produced comparable effects on body weight loss, food intake, and half-life in hTfR^het^ mice (Fig. S4A–C), confirming that placement of the TV35-containing chain on either the knob or hole arm did not significantly affect the pharmacokinetic or pharmacodynamic outcomes. Collectively, these results suggest that hTfR gene dosage does not affect the weight-loss efficacy of Fc(knob/hole-TV35)-FGF21.

**Figure 4.**
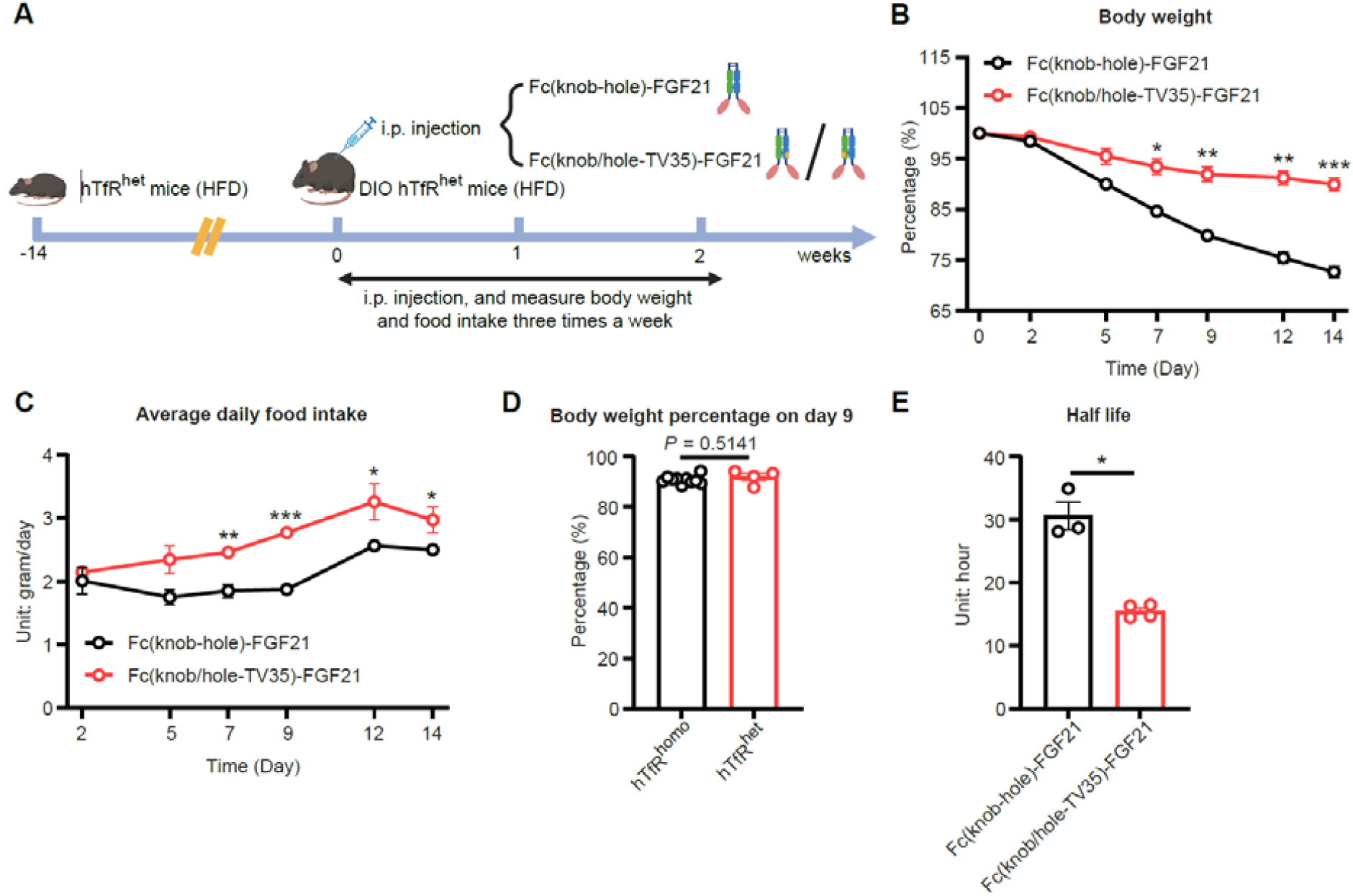
Reducing hTfR gene dosage did not improve the weight-loss efficacy of monovalent Fc(knob/hole-TV35)-FGF21. (A) Schematic diagram of the experimental design for hTfR^het^ mice treated with Fc(knob-hole)-FGF21 or Fc(knob/hole-TV35)-FGF21. (B) Relative body weight over time, presented as a percentage of body weight on day 0. DIO hTfR^het^ mice were treated with Fc(knob-hole)-FGF21 or Fc(knob/hole-TV35)-FGF21 three times per week for 2 weeks (n = 4 for the Fc(knob-hole)-FGF21 group; n = 4 for the Fc(knob/hole-TV35)-FGF21 group, including n = 2 for Fc(knob-TV35)-FGF21 and n = 2 for Fc(hole-TV35)-FGF21). (C) Corresponding average daily food intake per mouse, calculated over the specified intervals (days 0–2, 2–5, 5–7, 7–9, 9–12, and 12–14). (D) Body weight on day 9, expressed as a percentage of initial body weight, in hTfR^homo^ and hTfR^het^ mice treated with Fc(knob/hole-TV35)-FGF21 (n = 10 for the hTfR^homo^ group; n = 4 for the hTfR^het^ group). (E) Serum half-lives of Fc(knob-hole)-FGF21 and Fc(knob/hole-TV35)-FGF21 in hTfR^het^ mice (n = 4 for the Fc(knob-hole)-FGF21 group; n = 4 for the Fc(knob/hole-TV35)-FGF21 group, including n = 2 for Fc(knob-TV35)-FGF21 and n = 2 for Fc(hole-TV35)-FGF21). All data are presented as the mean ± s.e.m. \**p* < 0.05, \*\**p* < 0.01, and \*\*\**p* < 0.001, as determined using a mixed-effects two-way ANOVA or Student’s *t*-test, as appropriate.

## Discussion

The metabolic benefits of FGF21 involve coordinated actions in both the CNS [23, 24] and peripheral tissues [25, 26]. Given the sparse receptor expression in the brain [27], we hypothesized that engineering FGF21 to cross the BBB would enhance its body weight-lowering efficacy. Our data instead show that TV35-engineered Fc-FGF21 lost its weight-loss efficacy in hTfR^homo^ mice due to a severely shortened systemic half-life, indicating that sustained peripheral exposure is a major driver of the anti-obesity effect, consistent with clinical experience with long-acting FGF21 analogs. However, one limitation is that brain exposure was assessed using whole-brain homogenates at a single terminal time point; the present data do not establish BBB transcytosis or parenchymal delivery efficiency.

Our results demonstrate that increasing hTfR engagement from monovalent to bivalent formats progressively abolishes Fc–FGF21-mediated weight loss in hTfR^homo^ mice. This pattern is consistent with the current understanding that high-avidity TfR binding favors intracellular sequestration and lysosomal degradation in endothelial cells, whereas lower-avidity or monovalent engagement is more permissive for transcytosis but remains vulnerable to peripheral clearance in TfR-rich tissues. Recent work further indicates that efficient BBB transcytosis requires an intermediate range of TfR affinity that maximizes brain exposure while limiting lysosomal degradation and systemic receptor-mediated clearance [28]. Zygosity of hTfR mice seems to minimally affect the half-life and weight loss effects of engineered Fc-FGF21; this may be explained by abundant expression of TfR in brain capillary endothelial cells, reaching ∼10□–10□copies per cell [29] and rapid ligand-independent cycling of them [30], making zygosity-determined TfR expression differences in BBB endothelial cells no longer the limiting factor.

Given that TfR-based shuttles follow the normal recycling of TfR, they function as bidirectional, non-retentive conduits in which brain exposure is determined by the balance of influx, efflux, brain parenchymal retention, and peripheral sinks. For low-abundance CNS targets, such as the FGFR1c/β-Klotho receptor complex to which FGF21 binds, this low-retentive behavior is disadvantageous: in the absence of strong local retention, much of the dose is recycled back into the bloodstream and cleared by organs with high TfR expression. Within this framework, a TfR-based shuttle is most rational when the CNS target is abundant or broadly distributed, so that even transient exposure is sufficient to exert pharmacological effects.

CD98hc (encoded by gene *SLC3A2*) has emerged as an alternative BBB shuttle target with kinetic properties distinct from those of TfR. CD98hc-directed transport vehicles typically show slower but more sustained brain uptake, higher cumulative brain exposure, and longer parenchymal residence [31, 32]. Comparative studies indicate that TfR-targeted shuttles tend to yield rapid, high-peak brain signals followed by substantial cellular uptake and catabolism, whereas CD98hc-targeted shuttles provide slower uptake but more durable exposure and less intrinsic neuronal internalization [31-33]. For a payload such as FGF21, which exerts both central and peripheral effects on energy expenditure and body weight, a CD98hc-based shuttle might better support modest, sustained CNS exposure while preserving systemic pharmacokinetics.

Another potential design strategy would be a “releasable” construct in which a BBB-targeting Fc or antibody domain ferries FGF21 across the endothelium, followed by controlled dissociation or cleavage that releases FGF21 into the interstitial space, thereby decoupling CNS residence and peripheral clearance from continued TfR engagement. This could conceptually be achieved using protease-cleavable linkers that exploit CNS-enriched proteases, or affinity switches that promote release in abluminal compartments while remaining stable in circulation. However, our data suggest that any successful brain-targeted FGF21 design must preserve robust systemic exposure and treat CNS delivery as an additive, rather than exclusive, mechanism of action.

## Conclusions

In summary, our findings underscore that maintaining robust systemic exposure while minimizing high-avidity TfR engagement is essential for FGF21, and they support alternative brain-shuttle designs that are better suited to low-abundance CNS targets.

## Abbreviation list

Abbreviation: Definition
FGF21: Fibroblast Growth Factor 21
BBB: Blood–Brain Barrier
CNS: Central Nervous System
TfR / TfR1: Transferrin Receptor/Transferrin Receptor 1 hTfR / hTfR1 Human Transferrin Receptor 1
mTfR / mTfR1: Mouse Transferrin Receptor 1 FGFR1c Fibroblast Growth Factor Receptor 1c
TFRC: Transferrin Receptor Gene
CD98hc: CD98 Heavy Chain (SLC3A2)
Fc: Fragment Crystallizable (of IgG)
TV: Transport Vehicle
TV35: Transport Vehicle Variant 35
IgG1: Immunoglobulin G1
DIO: Diet-Induced Obese
WT: Wild-Type
HFD: High-Fat Diet
hTfR^homo^: Homozygous Humanized TfR
hTfR^het^: Heterozygous Humanized TfR

## Declarations

### Ethics approval and consent to participate

All animal protocols were approved by the Institutional Animal Care and Use Committee of Baylor College of Medicine. All experimental procedures complied with relevant ethical guidelines for animal research.

### Consent for publication

Not applicable.

### Availability of data and materials

The datasets generated and/or analyzed during the current study are available from the corresponding author on reasonable request.

### Author contributions

Y.Z., H.Z., and Y.D. designed the study. H.Z., Y.D., and Y.Z. wrote the manuscript. H.L., J.M., L.G.S., and S.Z. reviewed and edited the paper. H.Z., Y.D., H.L., and J.M. performed the experiments and/or analyzed the data. H.Z. and Y.D. prepared the figures. Y.Z. acquired funding for the study. All authors read and approved the final version of the manuscript.

