## Supplementary Information for "Transferrin receptor binding valency determines the circulating half-life and weight-loss efficacy of Fc-FGF21 in TfR-humanized mice"

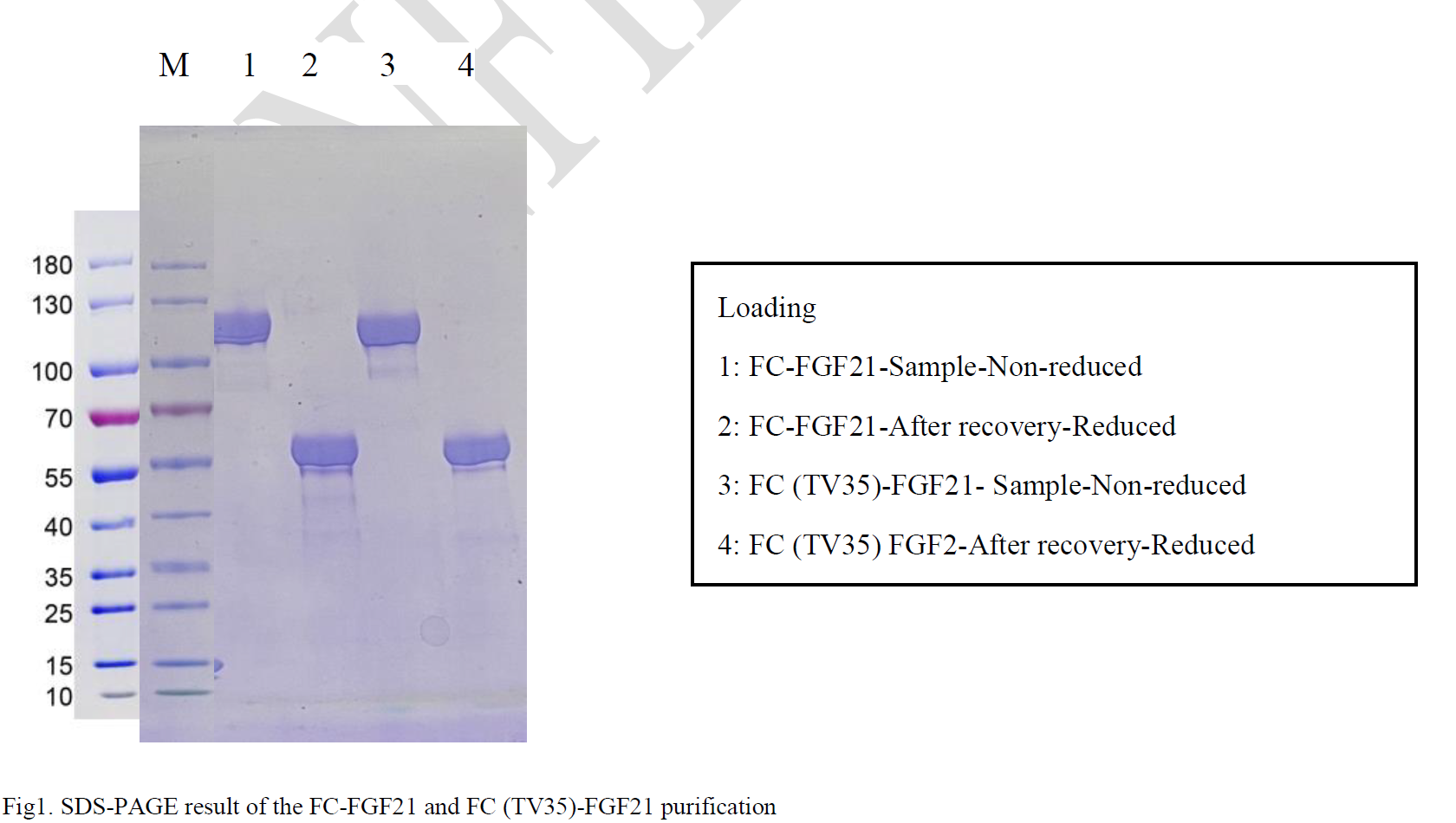

**Figure S1: SDS-PAGE gel image of protein purification, as shown in the vendor's certificate of analysis (COA).**

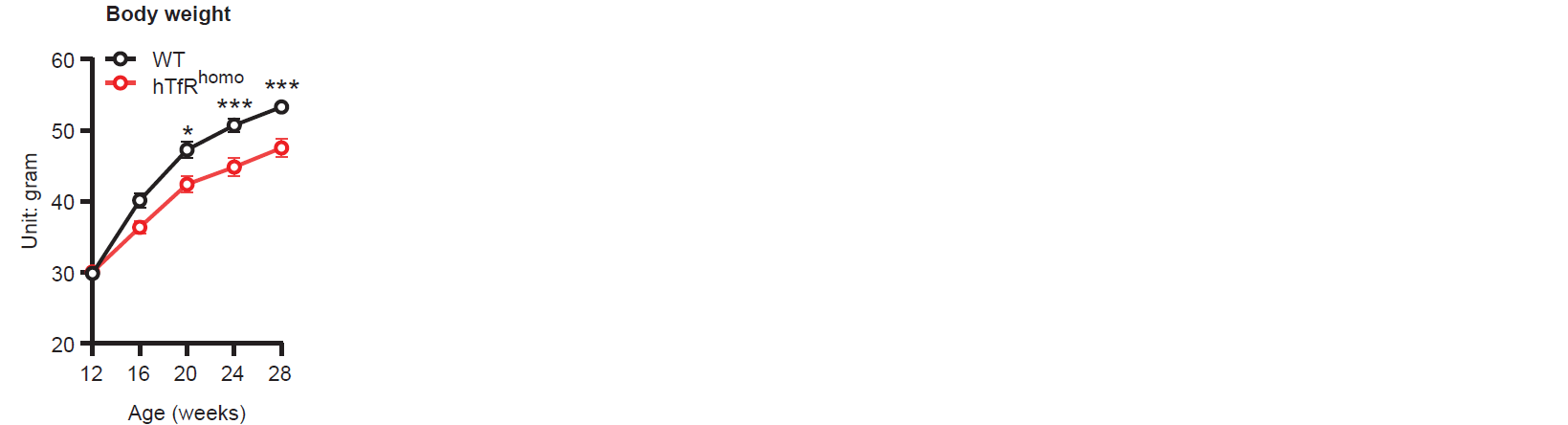

**Figure S2:** Body-weight trajectories of WT C57BL/6J (n = 20) and hTfR^homo^ mice (n = 21) on HFD, from 12 to 28 weeks of age. All data are presented as the mean ± s.e.m. **p* < 0.05, ***p* < 0.01, and ****p* < 0.001, as determined using a mixed-effects two-way ANOVA or Student’s *t*-test, as appropriate.

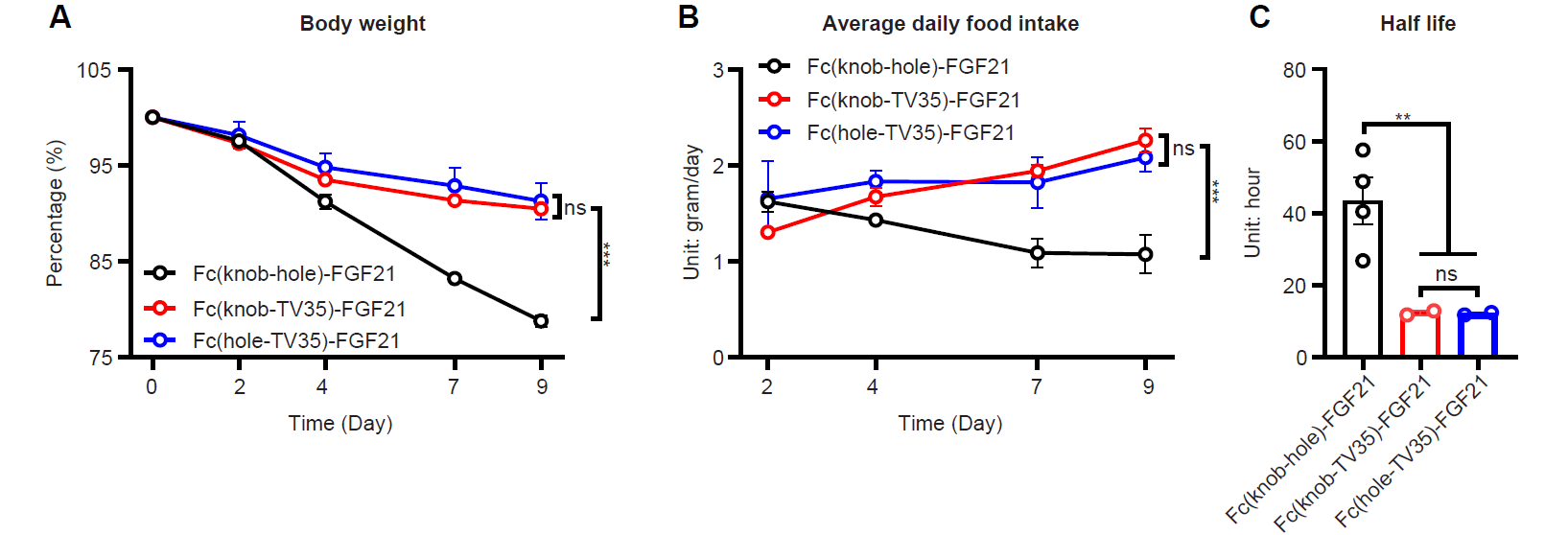

**Figure S3:** **The TV35 position on the Fc chain of Fc(knob/hole-TV35)-FGF21 does not affect its efficacy in DIO hTfR^homo^ mice.** (A) Relative body weight of DIO hTfR^homo^ mice changes over time, presented as a percentage of initial body weight on Day 0. DIO hTfR^homo^ mice were treated with Fc(knob-hole)-FGF21, Fc(knob-TV35)-FGF21 and Fc(hole-TV35)-FGF21 three times a week (n = 7 for the Fc(knob-hole)-FGF21 group, n = 5 for the Fc(knob-TV35)-FGF21 group and n = 5 for the Fc(hole-TV35)-FGF21 group). (B) Corresponding average daily food intake per mouse calculated over the specified intervals (Days 0–2, 2–4, 4–7, and 7–9). (C) Half-life of Fc(knob-hole)-FGF21, Fc(knob-TV35)-FGF21 and Fc(hole-TV35)-FGF21 in hTfR^homo^ mice (n = 4 for the Fc(knob-hole)-FGF21 group, n = 2 for the Fc(knob-TV35)-FGF21 group and n = 2 for the Fc(hole-TV35)-FGF21 group). All data are presented as mean ± s.e.m. ***p* < 0.01 and ****p* < 0.001, as determined by a mixed-effects two-way ANOVA and Student's *t-*test, as appropriate. ns: not significant.

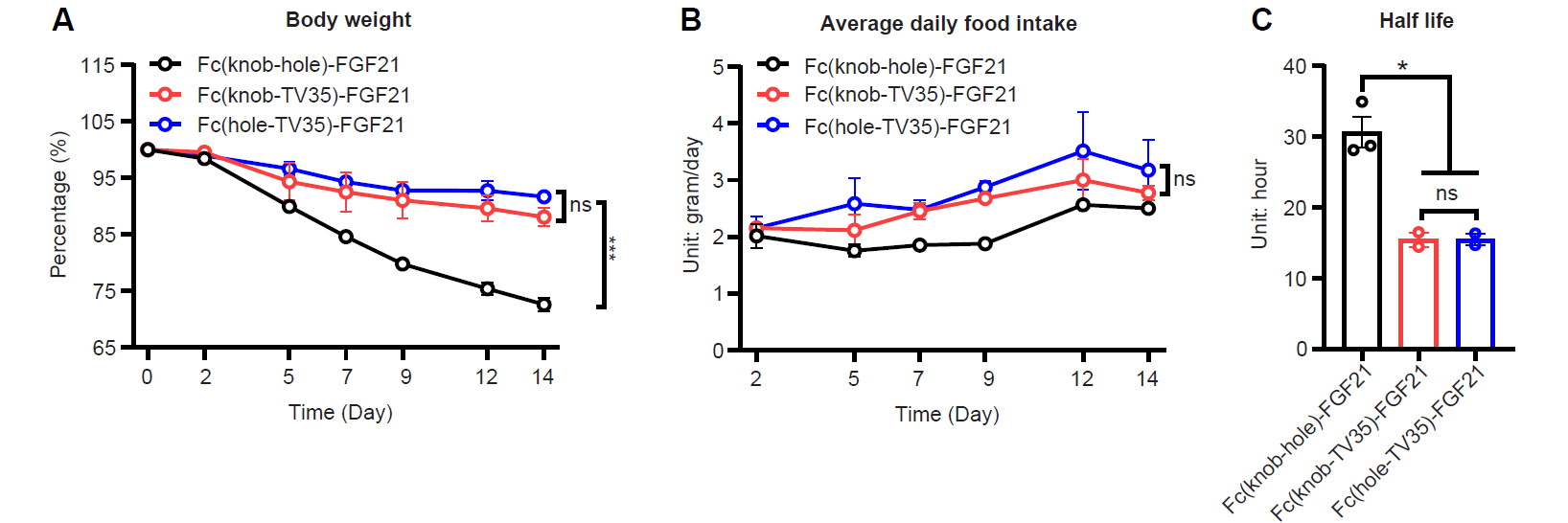

**Figure S4: The TV35 position on the Fc chain of Fc(knob/hole-TV35)-FGF21 does not affect its efficacy in DIO hTfR^het^ mice.** (A) Relative body weight of DIO hTfR^het^ mice changes over time, presented as a percentage of initial body weight on Day 0. DIO hTfR^het^ mice were treated with Fc(knob-hole)-FGF21, Fc(knob-TV35)-FGF21 and Fc(hole-TV35)-FGF21 three times a week (n = 4 for the Fc(knob-hole)-FGF21 group, n = 2 for the Fc(knob-TV35)-FGF21 group and n = 2 for the Fc(hole-TV35)-FGF21 group). (B) Corresponding average daily food intake per mouse calculated over the specified intervals (Days 0–2, 2–5, 5–7, 7–9, 9–12 and 12–14). (E) Half-life of Fc(knob-hole)-FGF21, Fc(knob-TV35)-FGF21 and Fc(hole-TV35)-FGF21 in hTfR^het^ mice (n = 4 for the Fc(knob-hole)-FGF21 group, n = 2 for the Fc(knob-TV35)-FGF21 group and n = 2 for the Fc(hole-TV35)-FGF21 group). All data are presented as mean ± s.e.m. **p* < 0.05 and ****p* < 0.001, as determined by a mixed-effects two-way ANOVA and Student's *t-*test, as appropriate. ns: not significant.
